# Microdeletions excluding *YWHAE* and *PAFAH1B1* cause a unique leukoencephalopathy and hypermobility syndrome: Further delineation of the 17p13.3 microdeletion spectrum

**DOI:** 10.1101/286146

**Authors:** Lisa T. Emrick, Jill A. Rosenfeld, Seema R. Lalani, Mahim Jain, Jill V. Hunter, Honey Nagakura, LaDonna L. Immken, Lindsay C. Burrage, Carlos A. Bacino, John W. Belmont, Brendan Lee

**Author notes:** Corresponding author: Lisa T. Emrick, MD. Kennedy Krieger Institute & Johns Hopkins School of Medicine, Baltimore, MD. Members of the Undiagnosed Diseases Network are collaborators; complete list provided as a supplement.

## Abstract

**Purpose:** Brain malformations are caused by 17p13.3 deletions: lissencephaly with deletions of the larger Miller-Dieker syndrome region or smaller deletions of only *PAFAH1B1*, as well as various abnormalities, including white matter changes, in the distinct syndrome due to deletions including *YWHAE* and *CRK* but sparing *PAFAH1B1.* We sought to understand the significance of 17p13.3 deletions between the *YWHAE*/*CRK* and *PAFAH1B1* loci.

**Methods:** We analyzed the clinical features of five individuals from four families with 17p13.3 deletions between and not including *YWHAE*/*CRK* and *PAFAH1B1* identified among individuals undergoing clinical chromosomal microarray testing.

**Results:** Four individuals from three families have multi-focal white matter lesions and hypermobile joints, while a fifth had a normal MRI. A combination of our patients and a review of those in the literature with white matter changes and deletions in this chromosomal region narrows the overlapping region for this brain phenotype to ∼345 kb, including 11 RefSeq genes, with *RTN4RL1* haploinsufficiency as the best candidate for causing this.

**Conclusion:** While previous literature has hypothesized dysmorphic features and white matter changes related to *YWHAE*, our cohort contributes evidence to the presence of additional genes within 17p13.3 required for proper brain development.

## Introduction

There is a spectrum of microdeletion syndromes associated with 17p13.3 deletions. Miller-Dieker syndrome (OMIM#247200) occurs when a deletion includes *YWHAE* and *PAFAH1B1* (*LIS1*). Isolated lissencephaly sequence (OMIM#6074332) occurs with smaller deletions involving *PAFAH1B1* but not *YWHAE*.^1^ Individuals with smaller deletions including *YWHAE* and *CRK* but sparing *LIS1* have growth restriction, cognitive impairment, dysmorphic features, and various brain abnormalities.^2-5^ White matter abnormalities have been reported in some patients with 17p13.3 microdeletions, and previous literature has hypothesized that this may be related to *YWHAE* haploinsufficiency.^2-4, 6^ We report a cohort of children who present with primarily white matter changes on brain MRI with mild dysmorphic features but no growth restriction or significant cognitive impairment with small interstitial 17p13.3 microdeletions. The patients have a primary static leukoencephalopathy with hypermobile joints but no evidence of developmental regression. The deletions are all proximal to *CRK*, which has been hypothesized to be associated with short stature,^2,3^ and distal to *PAFAH1B1*, associated with lissencephaly.^1^

## Materials & Methods

### Patient ascertainment

Patient 1 enrolled in the Undiagnosed Diseases Network (UDN) with a known chromosomal deletion involving 17p13.3, since the pathogenicity of the deletion and the etiology of her disease was not well established. Review of Baylor Genetics cytogenetic database for patients with similar deletions limited to the region between *YWHAE*/*CRK* and *PAFAH1B1* identified three other patients with white matter changes and one patient with a normal brain MRI. Two of these patients were brothers (Patients 2-3) and also enrolled in the UDN. A literature search with 17p13 search terms was used to identify other reports of similar white matter changes with deletions involving this region. The DECIPHER database was also searched for similar patients with deletions in the same region, although no individuals were identified with reported similar white matter changes.

### Chromosomal microarray testing

The patients were studied by either V8, V9, or V10 chromosomal microarrays (CMAs) designed by Baylor Medical Genetics Laboratories and manufactured by Agilent (Santa Clara, CA, USA). Patients 2, 4, and 5 were studied by the V8 array which included approximately 180,000 interrogating oligonucleotides, selected from Agilent’s online library (eArray; https://earray.chem.agilent.com/earray/). About 1,714 genes were covered on this array with an average of 4.2 probes per exon, excluding low-copy repeats and other repetitive sequences.^7^ Patients 3 and 1 were studied by the V9 and V10 array respectively, which targeted over 4,900 genes at the exon level plus 60,000 probes used for SNP analysis for the detection of uniparental disomy and absence of heterozygosity.^8^

### Whole exome sequencing

Clinical whole exome sequencing (WES) was performed in Patients 1 and 2 according to previously described methods.^9^

## Results

Patient 1 was born full-term via vaginal delivery to an 18-year-old mother with a pregnancy complicated by recurrent urinary tract infections. She weighed 3.03kg (33%) and required phototherapy for hyperbilirubinemia. She was discharged home at 4 days of life. She met all of her early milestones and attends regular classes but has difficulty with short-term memory.

At age 7 years she had recurrent vomiting after a viral illness. Additional symptoms included headaches, dizziness, inability to concentrate, and tingling in the legs. Her work-up revealed delayed gastric emptying and a hyperactive bladder. Brain and spine MRI showed significant white matter hyperintensities in the brain (Figure 1A), a Chiari I malformation, and a tethered cord. Her urinary and GI symptoms improved after cord release surgery. The white matter lesions remained unchanged on subsequent MRIs. Additional anomalies include a patent ductus arteriosus, a low IgA screen for celiac disease, and joint laxity (Beighton score 4/9). On examination she is normocephalic and nondysmorphic. Genetic workup included normal metabolic testing for arylsulfatase A, galactocerebrosidase, and thymidine levels. CSF analysis showed normal protein, glucose, and lactate. Infectious workup for tuberculosis, enterovirus and HSV was negative. CMA showed a 0.7 Mb *de novo* 17p13.3 deletion, between and not including *YWHAE* and *PAFAH1B1* (Figure 1D, Table 1). WES did not show any pathogenic variants or candidate genes associated with white matter changes or hypermobility, nor rare variants in the non-deleted 17p13.3 alleles.

**Table 1.**
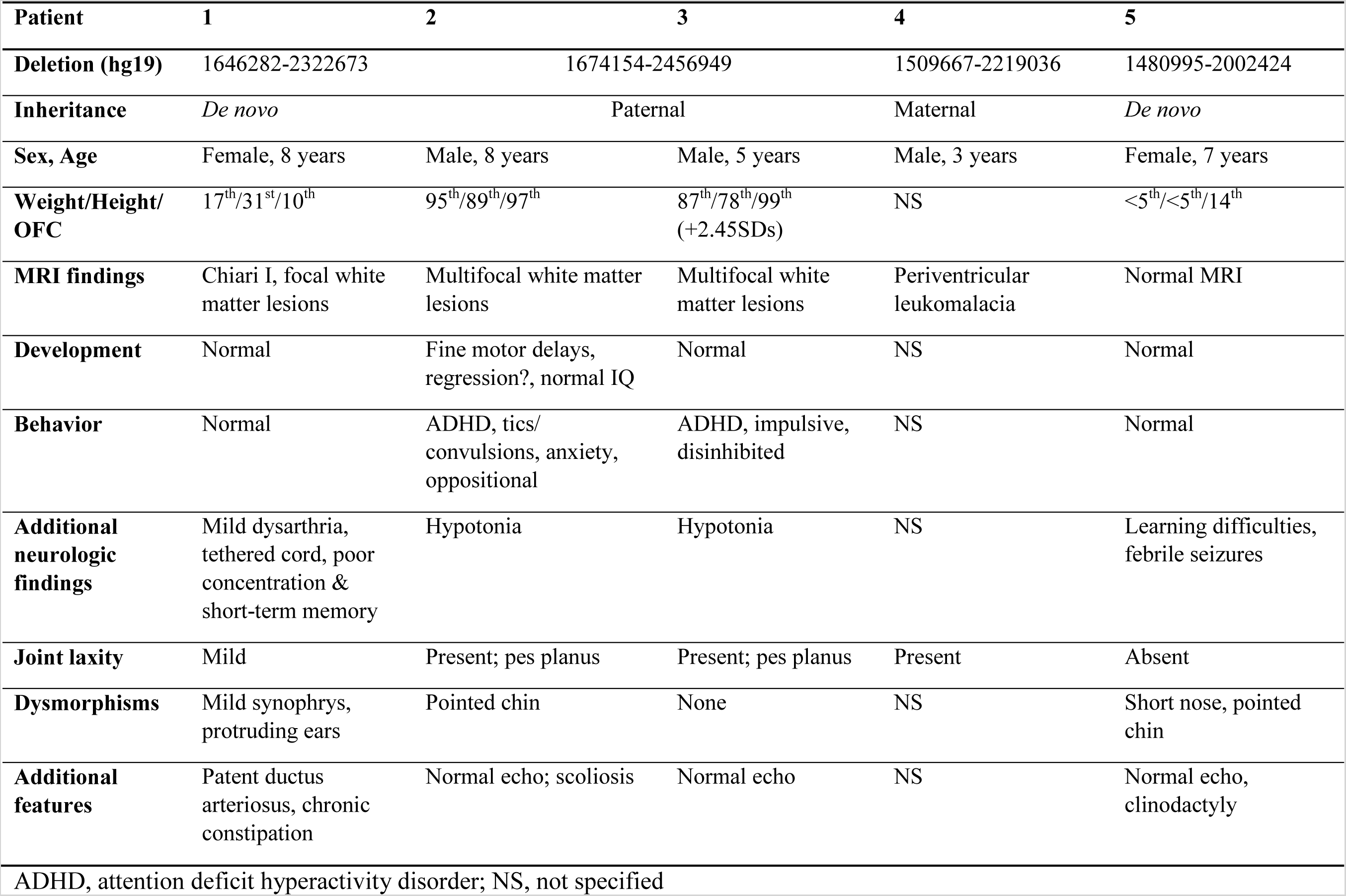
Genotypes and phenotypes in individuals with 17p13.3 deletions between *YWHAE*/*CRK* and *PAFAH1B1.*

**Figure 1.**
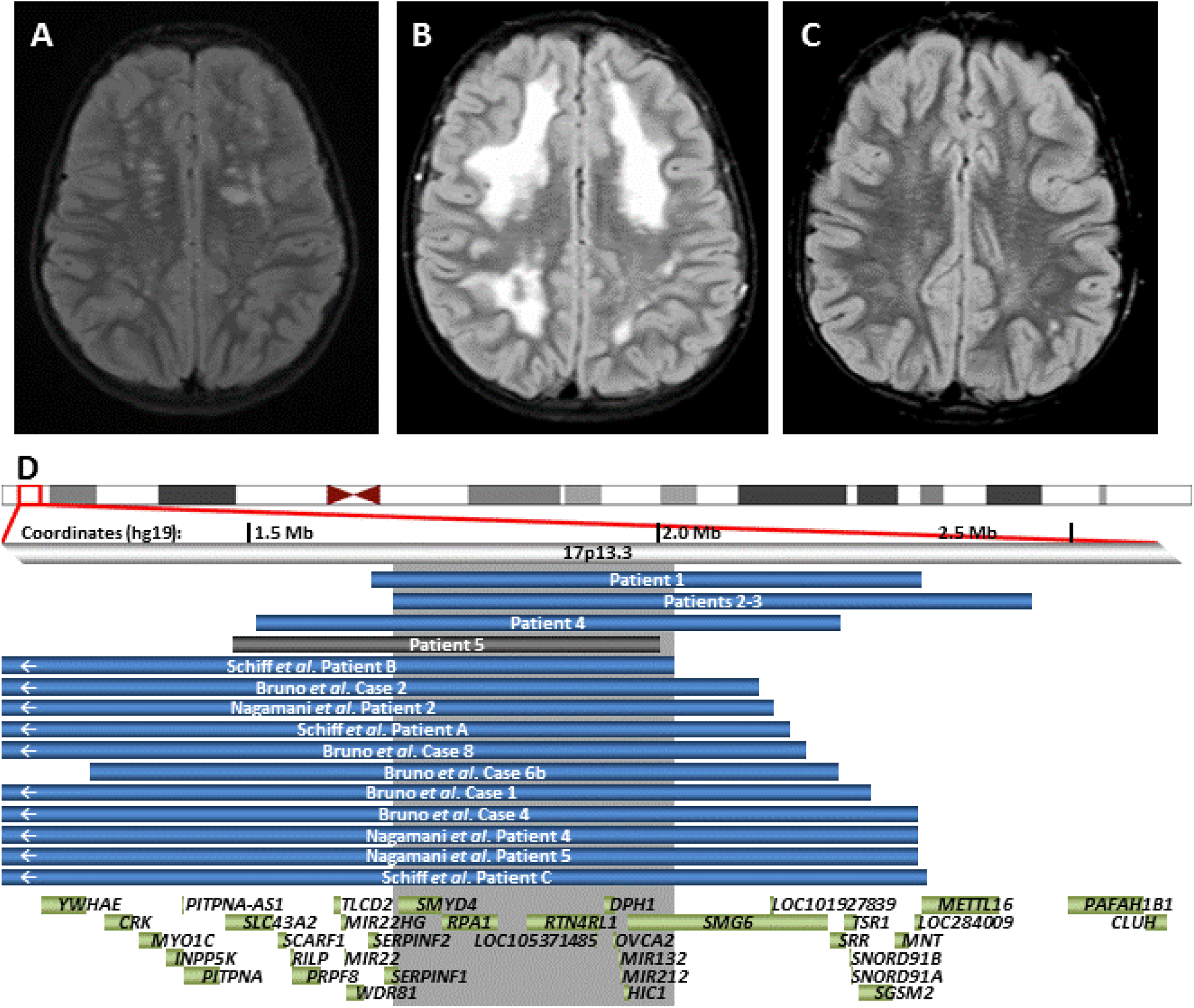
Leukoencephalopathies in patients with 17p13.3 deletions. Axial T2 weighted FLAIR (fluid attenuated inversion recovery) images demonstrate bilateral T2 hyperintensities in Patient 1 **(A)**, Patient 2 **(B)**, and Patient 3 **(C). (D)** Deletions associated with abnormal MRIs in our cohort and review of the literature are blue; the patient with a normal MRI is shown in gray. Gray shaded area represents the smallest region of overlap among cases with characteristic leukoencephalopathy. Genes in the region are shown as green boxes.

Patient 2 was born with a precipitous delivery at 33 weeks secondary to preterm labor to a 25-year-old mother. The pregnancy was complicated with vaginal bleeding in first and second trimesters and maternal anemia not requiring transfusions. He weighed 1.98kg (25%). He was in the neonatal intensive care unit for 24 days and had a grade I intraventricular hemorrhage. He had mild developmental delays and macrocephaly with head circumference >95%. At 6 years, he had possible developmental regression and worsening behaviors, and brain MRI showed extensive bilateral multifocal areas of T2 hyperintensities (Figure 1B). Multiple subsequent brain MRIs and MR spectroscopy showed no progression of these white matter changes. He has a normal IQ (100, at age 6) and was diagnosed with ADHD at age 8 with no additional concerns for regression. Physical exam showed macrocephaly with frontal bossing, hypermobile joints (Beighton score 6/9), and progressive pes planus. There was concern for connective tissue disorder, but echocardiogram and eye exam were normal. Genetic workup included normal urine mucopolysaccharide screening, serum very long chain fatty acids and arylsulfatase activity. CMA showed a 0.8 Mb 17p13.3 deletion (Figure 1D, Table 1) and a small deletion on 15q11.2 including only non-coding exons of *SNRPN*. The 17p13.3 deletion was inherited from his father and shared with a younger brother (Patient 3). Their father may also be similarly affected. He had problems with rolling his ankles when he was younger, as well as behavior issues. He has a cranial metal plate after a motor vehicle accident and therefore is unable to obtain a brain MRI, but a CT scan at the time of the accident showed white matter changes; additional details are not available. In Patient 2, WES did not show any pathogenic variants or candidate genes associated with white matter changes or hypermobility, nor rare variants in the non-deleted 17p13.3 alleles.

Patient 3 was born full-term to a 28-year-old mother with normal delivery and newborn course. He weighed 2.95kg (20%). He met all of his early milestones. Brain MRI at 3 years, performed due to macrocephaly and his brother’s MRI findings, showed patchy punctate bilateral multifocal areas of T2 hyperintensities (Figure 1C) and 2 small arachnoid cysts. Multiple subsequent brain MRIs and MR spectroscopy showed no progression of the white matter changes. His exam is significant for head circumference >98% with frontal bossing, hypermobile joints (Beighton score 6/9), and progressive pes planus. Echocardiogram showed mild mitral and tricuspid regurgitation. Ophthalmology noted mild hyperopia not requiring any intervention. Hyperkinetic behaviors were noted at 3 to 4 years old, and he was diagnosed with ADHD at age 5. Sleep study showed both mild central and obstructive sleep apnea and decreased time in REM sleep.

Patient 4 is a 3-year-old male. MRI showed periventricular leukomalacia. CMA showed a 0.7 Mb maternally inherited 17p13.3 deletion (Figure 1D, Table 1).

Patient 5 is a 7-year-old female referred to genetics for short stature. She has relative microcephaly with dysmorphic features including deep-set eyes, upslanting palpebral fissures, mild midface hypoplasia, and a pointed chin. Normal laboratory workup included thyroid studies, basic metabolic panel, urinalysis, IGF-1, and IGF binding protein. Sleep study at 6 years old demonstrated central sleep apnea. Brain MRI performed at 9 years did not show any white matter abnormalities. CMA showed a 0.5 Mb *de novo* 17p13.3 deletion (Figure 1D, Table 1).

## Discussion

Microdeletion syndromes involving 17p13.3 have a range of phenotypes, with the most severe being Miller-Dieker syndrome, with primary lissencephaly, growth failure, intellectual disability and dysmorphic features.^1^ Genotype-phenotype correlations have elucidated the roles of genes: *PAFAH1B1* involved in brain development^1^ and *CRK* in short stature.^2,3^ White matter changes have also been reported in some individuals with microdeletions in this region (Figure 1D, Table S1).^2-4^ We report a small cohort of children with white matter lesions, normal cognition and hypermobile joints with small 17p13.3 microdeletions between and not including *YWHAE*/*CRK* and *PAFAH1B1*. Previous literature has speculated a possible role for *YWHAE* in the brain anomalies, including the white matter changes.^2-4,6^ Our cohort suggests a different gene is likely responsible for the white matter findings, although it is likely that multiple 17p13.3 genes contribute to brain development. Due to the normal MRI in patient 5, whose deletion includes all genes in the smallest region of overlap (SRO) for this phenotype (Figure 1D), reduced penetrance is likely for these white matter changes.

White matter changes in the brain are often concerning for possible neurodegenerative leukodystrophy. Our cohort demonstrates that white matter changes can be associated with static, less severe conditions. The lesions in Patients 1-3 appear to be static based on multiple brain MRIs in each patient and the likely presence of similar findings in the father of Patients 2 and 3. Similar white matter changes have been described in individuals with other copy number variants (CNVs) that may contribute to their brain changes, although for the most part no causative genes have been confirmed within CNVs.^10^ Exceptions include haploinsufficiency of *MBP* (myelin basic protein) in 18q23 deletions and truncating deletions of *TM4SF20*, which have been linked to white matter changes on MRI.^11,12^

There are eleven RefSeq genes, *SERPINF1, SMYD4, RPA1, RTN4RL1, LOC105371485, DPH1, OVCA2, MIR132, MIR212, HIC1*, and *SMG6*, within our newly defined SRO (Figure 1D) for these white matter changes, and none has been previously associated with such brain findings. Most of these genes are not associated with Mendelian disorders, although *RTN4RL1, SMG6, MIR132* and *MIR212* are expressed in the brain. *SMG6* and *RTN4RL1* have the highest pLIs (probabilities of loss-of-function intolerance,^13^ 1.00 and 0.79, respectively) of the SRO genes. *RTN4RL1* regulates axonal and dendritic growth,^14^ and it may serve as a receptor for Nogo-66, a myelin-associated inhibitor.^15^ Of note, the Nogo-66 receptor gene *RTN4R* is located in 22q11.21, recurrent deletions of which can cause white matter abnormalities, a phenotype which has been hypothesized to be further modulated by polymorphisms in the nondeleted *RTN4R* allele.^16^ *SMG6* is involved in mRNA decay.^17^ As microRNAs *MIR132* and *MIR212* are noncoding, pLI scores are not applicable, although attenuation of *Mir132* in mice reduces neurite outgrowth.^18^

It has been hypothesized that *HIC1* haploinsufficiency (pLI score not available) contributes to the facial features and heart and gastrointestinal anomalies in Miller-Dieker syndrome,^19^ although these features are not prominent in our cohort. Given HIC1’s role in precartilaginous tissues and muscle development,^19^ this could be a candidate for the hypermobility in our patients. Additionally, as HIC1 acts as a tumor suppressor,^20^ future monitoring of patients with *HIC1* deletions for risk of neoplasia is possibly warranted. The remainder of the genes in the SRO have low pLI scores, and two (*SERPINF1* and *DPH1*) are associated with recessive disease (OMIM#613982 and #616901), so it is less likely that these genes are haploinsufficient. However, as these white matter changes may have minimal outward phenotypic impacts, we cannot rule out the possibility that one of these other genes with more tolerance to loss-of-function contributes to the MRI findings.

When white matter changes are found, long-term follow-up is recommended to evaluate whether the leukoencephalopathy is static. Given the likely presence of such changes in an adult (father of Patients 2-3) and normal development in our patients, some reassurance may be given to families with similar 17p13.3 deletions, but clinical follow-up is still recommended as there may be phenotypic variability. Additional factors, such as the prematurity in Patient 2, may increase susceptibility for the brain phenotype; this may also explain why the phenotype appears more prominent in him (Figure 1B).

Our cohort contributes evidence to the presence of multiple genes within 17p13.3 required for normal brain development. Identification of additional individuals with deletions and pathogenic variants within this region may help to identify specific genes responsible. Additional studies, including diffusion tensor imaging and downstream techniques such as RNAseq to evaluate for abnormal expression of genes within and adjacent to the deletion and in relevant developmental pathways, may assist with elucidating genetic etiologies of abnormal myelination.

## ACKNOWLEDGEMENTS

We thank the patients and their families for their participation in this research. Research reported in this manuscript was supported by the NIH Common Fund, through the Office of Strategic Coordination/Office of the NIH Director under Award Number U01 HG007709-01. The content is solely the responsibility of the authors and does not necessarily represent the official views of the National Institutes of Health. This work was also supported by the BCM Intellectual and Developmental Disabilities Research Center (HD024064) from the Eunice Kennedy Shriver National Institute of Child Health and NIGMS T32 GM007526 (BL, LE, MJ, LB, JB).

## Conflict of interest notification

The Department of Molecular and Human Genetics at Baylor College of Medicine receives revenue from clinical genetic testing conducted at Baylor Genetics, including chromosomal microarray testing. John W. Belmont is an employee of Illumina. Honey Nagakura is an employee of LabCorp. The remaining authors have no conflicts to declare.

---

**Table S1.** Reports of white matter abnormalities in individuals with 17p13.3 microdeletions.^**1-3**^

| Patient identifier | Brain MRI | Physical features | Deletion (hg19; inheritance) |
| --- | --- | --- | --- |
| **Bruno *et al.***<br>**Patient 1** | White matter, wide perivascular spaces, low cerebellar tonsils | Mild ID, bitemporal narrowing, high anterior hairline, laterally extended eyebrows, maxillary prominence, micrognathia | 1.07 Mb: 1188272–2256085 (paternal) |
| **Bruno *et al.***<br>**Patient 2** | White matter, low cerebellar tonsils | Moderate ID, frontal bossing, laterally extended eyebrows, broad nasal tip, maxillary prominence, retrognathia, 5<sup>th</sup> finger clinodactyly | 2.12 Mb: 514–2118914 (paternal) |
| **Bruno *et al.***<br>**Patient 4** | White matter – wide perivascular spaces | Mild ID, speech delay, midface retrusion, broad high anterior hairline, laterally extended eyebrows, maxillary prominence | 1.42 Mb: 895639–2311107 (*de novo*) |
| **Bruno *et al.***<br>**Patient 6b** | White matter | Developmental delay, broad nasal tip, maxillary prominence, 5<sup>th</sup> finger clinodactyly | 0.91 Mb: 1307777–2217389 (maternal) |
| **Bruno *et al.***<br>**Patient 8** | White matter, wide perivascular spaces, hypoplastic adenohypophysis and olfactory tracts | Moderate ID, triangular facies, frontal bossing, eyelide coloboma, maxillary prominence, micrognathia | 2.15 Mb: 29169–2177066 (*de novo*) |
| **Schiff *et al.***<br>**Patient A** | White matter, enlargement of subarachnoid spaces | Macrocephaly, mild to moderate developmental disorder, prominent forehead, broad nasal root | 1.93 Mb: 223182–2157982 (*de novo*) |
| **Schiff *et al.***<br>**Patient B** | White matter | Microcephaly, mild to moderate developmental delay, prominent forehead, downslanting fissures, epichanthus, low set ears, high arched palate, retrognathia | 1.79 Mb: 223182–2016759 (*de novo*) |
| **Schiff *et al.***<br>**Patient C** | White matter, enlarged frontal horns | Macrocephaly, mild to moderate developmental disorder, prominent forehead, broad nasal root, retrognathia | 1.42 Mb: 902315–2322673 (*de novo*) |
| **Nagamani *et al.***<br>**Patient 2** | White matter – subcortical, Chiari I | Growth restriction, mild delay, prominent forehead, tall vertex, broad nasal root, PDA and PFO and coloboma iris | 1.07 Mb: 1074030–2137299 (*de novo*) |
| **Nagamani *et al.***<br>**Patient 4** | Prominent Virchow Robin spaces, Chiari I | Growth restriction, mild delay, prominent forehead, tall vertex, broad nasal root, epicanthus inversus, retrusion of mandible, pectus excavatum, undescended testes | 1.64 Mb: 668533–2312597 (*de novo*) |
| **Nagamani *et al.***<br>**Patient 5** | Prominent Virchow Robin spaces, white matter – subcortical | Short stature, mild delay, prominent forehead, tall vertex, broad nasal root, PDA | 2.31 Mb: 1–2314108 (*de novo*) |

ID, intellectual disability; PDA, patent ductus arteriosus; PFO, patent foramen ovale

## REFERENCES

1. Dobyns WB, Das S. LIS1-Associated Lissencephaly/Subcortical Band Heterotopia. In: Adam MP, Ardinger HH, Pagon RA, et al., eds. GeneReviews((R)). Seattle (WA)1993.

2. Nagamani SC, Zhang F, Shchelochkov OA, et al. Microdeletions including *YWHAE* in the Miller-Dieker syndrome region on chromosome 17p13.3 result in facial dysmorphisms, growth restriction, and cognitive impairment. J Med Genet. Dec 2009;46(12):825–833.

3. Bruno DL, Anderlid BM, Lindstrand A, et al. Further molecular and clinical delineation of co-locating 17p13.3 microdeletions and microduplications that show distinctive phenotypes. J Med Genet. May 2010;47(5):299–311.

4. Schiff M, Delahaye A, Andrieux J, et al. Further delineation of the 17p13.3 microdeletion involving *YWHAE* but distal to *PAFAH1B1*: four additional patients. Eur J Med Genet. Sep-Oct 2010;53(5):303–308.

5. Tenney JR, Hopkin RJ, Schapiro MB. Deletion of 14-3-3{varepsilon} and *CRK*: a clinical syndrome with macrocephaly, developmental delay, and generalized epilepsy. J Child Neurol. Feb 2011;26(2):223–227.

6. Noor A, Bogatan S, Watkins N, Meschino WS, Stavropoulos DJ. Disruption of YWHAE gene at 17p13.3 causes learning disabilities and brain abnormalities. Clin Genet. Feb 2018;93(2):365–367.

7. Chaudhry A, Noor A, Degagne B, et al. Phenotypic spectrum associated with PTCHD1 deletions and truncating mutations includes intellectual disability and autism spectrum disorder. Clin Genet. Sep 2015;88(3):224–233.

8. Wiszniewska J, Bi W, Shaw C, et al. Combined array CGH plus SNP genome analyses in a single assay for optimized clinical testing. Eur J Hum Genet. Jan 2014;22(1):79–87.

9. Yang Y, Muzny DM, Xia F, et al. Molecular findings among patients referred for clinical whole-exome sequencing. JAMA. Nov 12 2014;312(18):1870–1879.

10. Garcia-Cazorla A, Sans A, Baquero M, et al. White matter alterations associated with chromosomal disorders. Dev Med Child Neurol. Mar 2004;46(3):148–153.

11. Loevner LA, Shapiro RM, Grossman RI, Overhauser J, Kamholz J. White matter changes associated with deletions of the long arm of chromosome 18 (18q-syndrome): a dysmyelinating disorder? AJNR Am J Neuroradiol. Nov-Dec 1996;17(10):1843–1848.

12. Wiszniewski W, Hunter JV, Hanchard NA, et al. TM4SF20 ancestral deletion and susceptibility to a pediatric disorder of early language delay and cerebral white matter hyperintensities. Am J Hum Genet. Aug 8 2013;93(2):197–210.

13. Lek M, Karczewski KJ, Minikel EV, et al. Analysis of protein-coding genetic variation in 60,706 humans. Nature. Aug 18 2016;536(7616):285–291.

14. Pignot V, Hein AE, Barske C, et al. Characterization of two novel proteins, NgRH1 and NgRH2, structurally and biochemically homologous to the Nogo-66 receptor. J Neurochem. May 2003;85(3):717–728.

15. Zhang L, Kuang X, Zhang J. Nogo receptor 3, a paralog of Nogo-66 receptor 1 (NgR1), may function as a NgR1 co-receptor for Nogo-66. J Genet Genomics. Nov 20 2011;38(11):515–523.

16. Perlstein MD, Chohan MR, Coman IL, et al. White matter abnormalities in 22q11.2 deletion syndrome: preliminary associations with the Nogo-66 receptor gene and symptoms of psychosis. Schizophr Res. Jan 2014;152(1):117–123.

17. Ottens F, Boehm V, Sibley CR, Ule J, Gehring NH. Transcript-specific characteristics determine the contribution of endo- and exonucleolytic decay pathways during the degradation of nonsense-mediated decay substrates. RNA. Aug 2017;23(8):1224–1236.

18. Vo N, Klein ME, Varlamova O, et al. A cAMP-response element binding protein-induced microRNA regulates neuronal morphogenesis. Proc Natl Acad Sci U S A. Nov 8 2005;102(45):16426–16431.

19. Grimm C, Sporle R, Schmid TE, et al. Isolation and embryonic expression of the novel mouse gene Hic1, the homologue of HIC1, a candidate gene for the Miller-Dieker syndrome. Hum Mol Genet. Apr 1999;8(4):697–710.

20. Fleuriel C, Touka M, Boulay G, Guerardel C, Rood BR, Leprince D. HIC1 (Hypermethylated in Cancer 1) epigenetic silencing in tumors. Int J Biochem Cell Biol. Jan 2009;41(1):26–33.

